# Genome wide CRISPR screen for *Pasteurella multocida* toxin (PMT) binding proteins reveals LDL Receptor Related Protein1 (LRP1) as crucial cellular receptor

**DOI:** 10.1101/2022.08.04.502755

**Authors:** Julian Schöllkopf, Thomas Müller, Lena Hippchen, Teresa Müller, Sonja Großmann, Raphael Reuten, Rolf Backofen, Norbert Klugbauer, Joachim Orth, Gudula Schmidt

## Abstract

PMT is a protein toxin produced by *Pasteurella multocida* serotypes A and D. As causative agent of atrophic rhinitis in swine, it leads to rapid degradation of the nasal turbinate bone. The toxin acts as a deamidase to modify a crucial glutamine in heterotrimeric G proteins, which results in constitutive activation of the G proteins and permanent stimulation of numerous downstream signaling pathways.

Using a lentiviral based genome wide CRISPR knockout screen in combination with a lethal toxin chimera, consisting of full length inactive PMT and the catalytic domain of diphtheria toxin, we identified the LRP1 gene encoding the Low-Density Lipoprotein Receptor-related protein 1 as a critical host factor for PMT function. Loss of LRP1 reduced PMT binding and abolished the cellular response and deamidation of heterotrimeric G proteins, confirming LRP1 to be crucial for PMT uptake. Expression of LRP1 or cluster 4 of LRP1, respectively, restored intoxication of the knockout cells. In summary our data demonstrate secretory cells as entry site of PMT into airway epithelia and present LRP1 as crucial host entry factor for PMT intoxication by acting as its primary cell surface receptor.

## Introduction

Pathogenic bacteria produce and release protein toxins which enter mammalian cells to dictate their appearance and behavior. Frequently, these toxins are enzymes which modify specific targets within the cells. Therefore, a lot of those proteins comprise everything needed for host cell binding and uptake into the cytosol: a binding domain interacting with a cell surface receptor to mediate endocytosis, secondly a membrane insertion domain involved in the passage of the catalytic domain through the endosomal membrane and third the catalytic domain. Such toxins are named AB toxins, because they encompass binding properties (B) and catalytic activity (A) in a single protein.

*Pasteurella multocida*, a gram-negative bacterium mostly involved in zoonotic diseases like pasteurellosis and atrophic rhinitis, produces the large (146 kDa) AB protein toxin Pasteurella multocida toxin (PMT) as major virulence factor. PMT is a deamidase which modifies a specific glutamine in the switch-II region of the alpha subunits of heterotrimeric G proteins (G_i_, G_q_ and G_12/13_) (1). This leads to direct and permanent activation and stimulation of numerous signaling pathways ending up in mitogenic signaling, reorganization of the cytoskeleton, de-differentiation and many other cellular responses (2). In pigs, *Pasteurella multocida* causes atrophic rhinitis with severe degradation of the turbinate bone (3). Therefore, the main target cells of PMT present cells involved in bone degradation and formation as well as the respiratory system. The toxin stimulates osteoclastogenesis and inhibits osteoblast differentiation (4). Moreover, osteocytes are activated to stimulate the differentiation of osteoclast precursor cells by increased secretion of the receptor activator of nuclear factor κB ligand (RANKL) (5). Following binding to a cellular receptor, PMT is taken up into mammalian cells by receptor-mediated endocytosis. It is then released into the cytosol from acidified endosomes. In the presence of Bafilomycin A1, an inhibitor of the endosomal proton pump (blocking acidification), intoxication by PMT is completely abolished (6). However, the exact mechanism how PMT crosses the endosomal membrane is not known. Two hydrophobic helices within the N-terminal binding and translocation domain (Fig.1) are proposed to be involved in toxin translocation. Like many protein toxins, PMT is composed of the typical functional domains of bacterial AB toxins. Receptor binding and translocation through the endosomal membrane is mediated by the N-terminal amino acids 1 to 575 followed by three C-terminal domains C1 to C3. The minimal region encompassing the deamidase activity (C3) is located at the very C-terminus of the toxin (residues 1106 to 1285) with the catalytic triad typically present in deamidases: Cys1165, His1205 and Asp1220. The C1 domain (residues 576 to720) mediates attachment to the inner surface of the cellular plasma-membrane, while the function of domain C2 (residues 721 to 1105) has not been elucidated (Fig. 1, S1 for review see (7, 8)). In this study we generated a fusion of catalytically inactive full length PMT (PMT-C1165S) and the catalytic domain of *Corynebacterium diphtheriae* toxin (DTa) (9), because PMT itself is not able to rapidly kill mammalian cells. Diphtheria toxin catalyzes the ADP-ribosylation of elongation factor thermo unstable (EF-Tu) which leads to its inactivation and therefore to the block of protein synthesis and eventually cell death. Ribosylation occurs at a diphthamide, a post-translationally modified histidine in EF-Tu (10, 11).

**Fig. 1:**
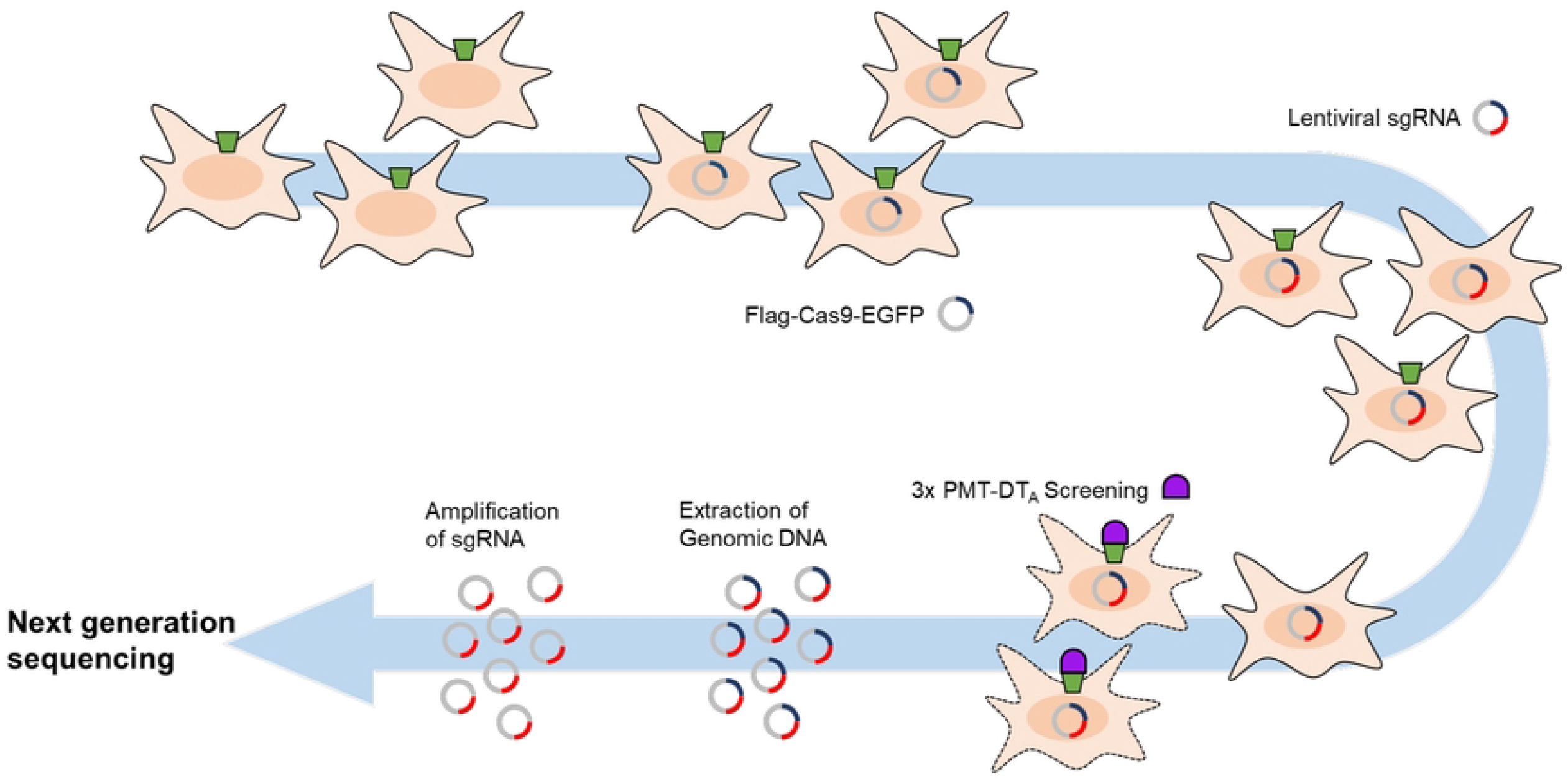
schematic presentation of the CRISPR/Cas9-Knock out screen. Mouse embryonic fibroblasts (MEF) stably expressing Flag-Cas9-EGFP were transduced with a lentiviral CRISPR-library. Cells were treated three times with PMT(C1165S)DTa and surviving cells grown. Genomic DNA was extracted, inserted sgRNA amplified and sequenced.

Using a CRISPR knockout library, we identified the LDL Receptor Related Protein1 (LRP1, CD91) as a cellular receptor crucial for PMT binding and uptake. LRP1 belongs to the family of low-density lipoprotein receptors which is encompassed of a single pass transmembrane protein composed of four extracellular domains (called cluster 1-4, LRP1α, 515 kDa) as well as a transmembrane and a cytoplasmic domain, which together form LRP1β (85 kDa). The two LRP parts originate from a single precursor-protein which is cleaved by furin within the endoplasmic reticulum (12). The chaperone receptor associated protein (RAP) escorts LRP1 from the endoplasmic reticulum to the Golgi apparatus. The tight binding of RAP to LRP1 has been used for competition experiments proving newly identified receptor ligands (13). On the cell surface the two parts remain non-covalently linked. More than 40 distinct ligands have been identified to interact with LRP1 including apolipoproteins and cholesteryl esters in remnants (14, 15), extracellular matrix (ECM) components such as fibronectin (16, 17), a minor-group common cold virus (18) and Rift Valley fever virus (19) as well as bacterial toxins like *Clostridium perfringens* TpeL (13), *Pseudomonas* Exotoxin A (ExoA, (20)), TcdA produced by *Clostridoides difficile* (21) and the vacuolating toxin (VacA) from *Helicobacter pylori* (22). These diverse ligands do not contain a common binding motif. Thus, LRP1 is suggested to be a general endocytosis receptor. Global deletion of LRP1 in mice is lethal. Therefore, conditional gene knockout has been used to elucidate its function in several tissues. In osteoblasts, knockout of LRP1 promotes osteoporosis (23) and reduces delivery of lipoproteins and lipophilic vitamins to bone (24). Moreover, several signaling pathways for example such induced by cytokines and growth factors are modulated by LRP1 (17, 25, 26).

## Results

### Identification of essential binding partners for PMT on mammalian cells

We established a CRISPR/Cas9 based knockout screen for identification of the cellular receptor for the bacterial toxin Pasteurella multocida toxin (PMT) mainly following the excellent work performed by the group of Y. Horiguchi (27). The method selects cells which are not killed by the toxin or the toxin chimera of interest because an essential protein for binding, uptake or modification of the target, respectively is missing due to gene knockout. However, PMT itself is not able to kill cells. Therefore, we generated a fusion toxin composed of the catalytically inactive mutant of PMT (PMT-C1165S) and the functional catalytic domain of diphtheria toxin (DTa) (Fig. S1). For an effective and target-oriented search for the cellular receptor, we established the following CRISPR/Cas9 based knockout screen: Two mouse cell lines were generated which stably express FLAG-Cas9-EGFP (27). These cells were transduced with a lentiviral CRISPR library (Add Gene, Kosuke Yusa) using a multiplicity of infection (MOI) of 0.3 to generate mainly single gene knockouts by avoiding multiple transduction. Cells were than treated with the PMT-DTa chimera. Non-transduced cells as well as cells with a functional receptor and uptake machinery most likely were killed by the toxin. Only cells with a receptor knockout or cells which block the toxins activity by other means should have survived. These cells were further amplified, genomic DNA extracted and sequences which encode the single guide RNA were amplified and analyzed by next generation sequencing (Eurofins). A schematic overview of the method is shown in Fig. 1.

The sequence data was analyzed for significantly recovered sgRNAs and their corresponding genes were identified. The results are presented in Fig. 2 and Tab. S2. Two genes (Dph 1 und Dph 5) encoding enzymes required for diphthamide biosynthesis were present within the first ten hits. Diphthamide is a histidine which is post translationally modified in some proteins. Since diphtheria toxin selectively modifies a diphthamide present in elongation factor 2, the enzymes identified are crucial for the toxins action. Therefore, these targets proof to be a reasonable readout of our CRISPR screen. The top hit in the screen and the only transmembrane cell surface protein was the Low-Density Lipoprotein Receptor-related protein 1 (LRP1) (Tab. S2). LRP1 belongs to the family of low-density lipoprotein receptors and is highly expressed in many tissues, including the respiratory system (proteinatlas.org), an important entrance for *Pasteurella multocida*. The LRP1 protein is composed of four extracellular domains (cluster 1-4), a transmembrane and a cytoplasmic domain.

**Fig. 2:**
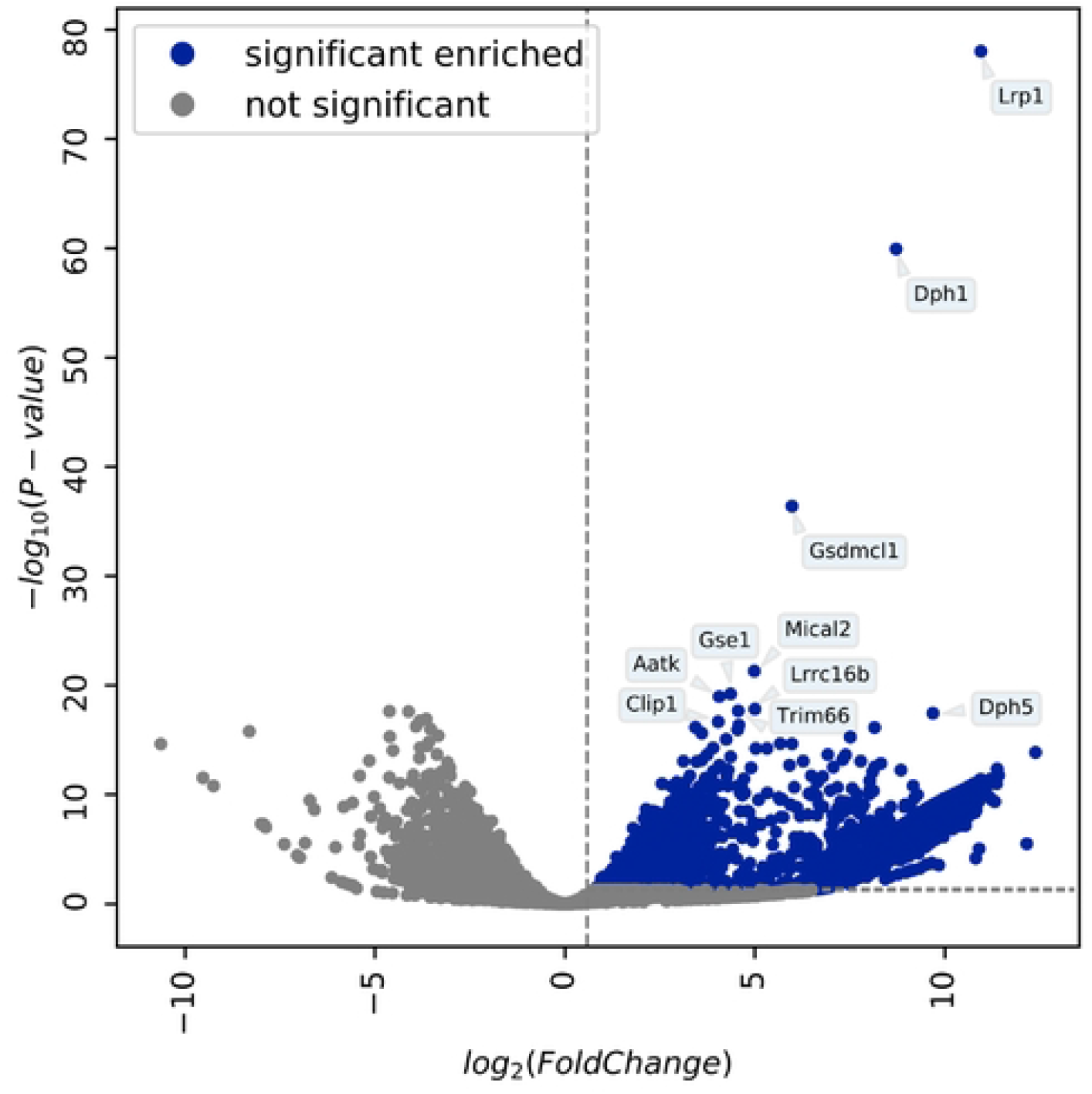
volcano plot showing significantly enriched genes. The volcano plot shows gene expression changes between cells treated with lethal PMT-DTa chimera and those only transduced with the CRISPR library. All reads were mapped locally using BWA-MEM [5,6], then quantified with featureCounts [4], and finally fold changes between the condition were calculated by DESeq2 [7] (see galaxy history). The volcano plot was drawn with the bioinfokit toolkit [8]. Significantly enriched genes, having a positive fold change above 0.584 and a p-value lower or equal 5%, are shown in blue. The significance thresholds are marked by gray-dotted lines. The ten most significant genes are highlighted with their name. Non-significant genes are colored in gray.

### LRP1 knockout prohibits binding and uptake of PMT

To analyze whether LRP1 is crucial for binding and uptake of PMT, we used mouse embryonic fibroblasts (MEF) deficient for the LRP1 gene (12).

First, we studied viability of cells treated with the fusion toxin PMT-DTa in a concentration-dependent manner. In case LRP1 would be crucial for toxin binding and/or uptake, the knockout cells should survive. In Fig. 3a cell viability was plotted in % of living wildtype fibroblasts and LRP1 knockout fibroblasts (LRP1^-/-^) normalized to the respective untreated control with increasing concentration of PMT-DTa. Whereas the wildtype MEFs are killed already at low concentrations of PMT-DTa, there was no effect on the LRP1 knockout cells even at high concentrations. This indicates that LRP1 is crucial for binding/uptake of the fusion toxin. To study whether this holds true for PMT without DTa, we analyzed the binding properties of Alexa488 labeled PMT (PMT_488_) to each cell line by FACS. As shown in Fig. 3b, binding of PMT_488_ to both cell lines increased in a concentration-dependent manner. However, fluorescence intensity of wildtype cells was much higher compared to LRP1^-/-^ cells. This indicates that LRP1 is involved in binding of PMT to mammalian cells. To study whether the binding of PMT_488_ to LRP1 is crucial for intoxication, we directly studied PMT-induced intoxication of MEF wildtype and LRP1 knockout cells. Therefore, we made use of an antibody which detects deamidated Gα (Gα Q209E). Wildtype and knockout MEF were treated with PMT for 2 to 6 h or left untreated. Cells were lysed and the lysates analyzed for deamidated Gα, the presence of LRP1 and tubulin as loading control by Western-blotting as indicated. As shown in Fig. 3c (top), the G proteins in the lysates of wildtype MEF were already deamidated 2 h following toxin treatment, whereas there was no deamidation detectable in the lysates of LRP1^-/-^ cells up to 6 h after PMT exposure. The data indicate that LRP1 is crucial for uptake of PMT into cells and thus for intoxication. To further strengthen the finding, we made use of a known chaperone and ligand of LRP1, receptor-associated protein (RAP) to compete with PMT for LRP1 binding (Martin et al, 2008). Therefore, MEF cells were incubated with buffer, 1 nM or 10 nM of PMT, respectively in the presence or absence of recombinant glutathione-S-transferase (GST) or GST-RAP (1 µM each), incubated for 2h, washed and lysed. Lysates were then analyzed for deamidated Gα. As shown in Fig. 3c (bottom), deamidation of Gα in PMT-treated cells was significantly reduced when the cells were co-incubated with GST-RAP, but not in toxin-treated cells co-treated with GST. This indicates that GST-RAP but not GST interfered with the binding/action of PMT, suggesting a crucial role of the RAP receptor LRP1 for intoxication with PMT.

**Fig.3:**
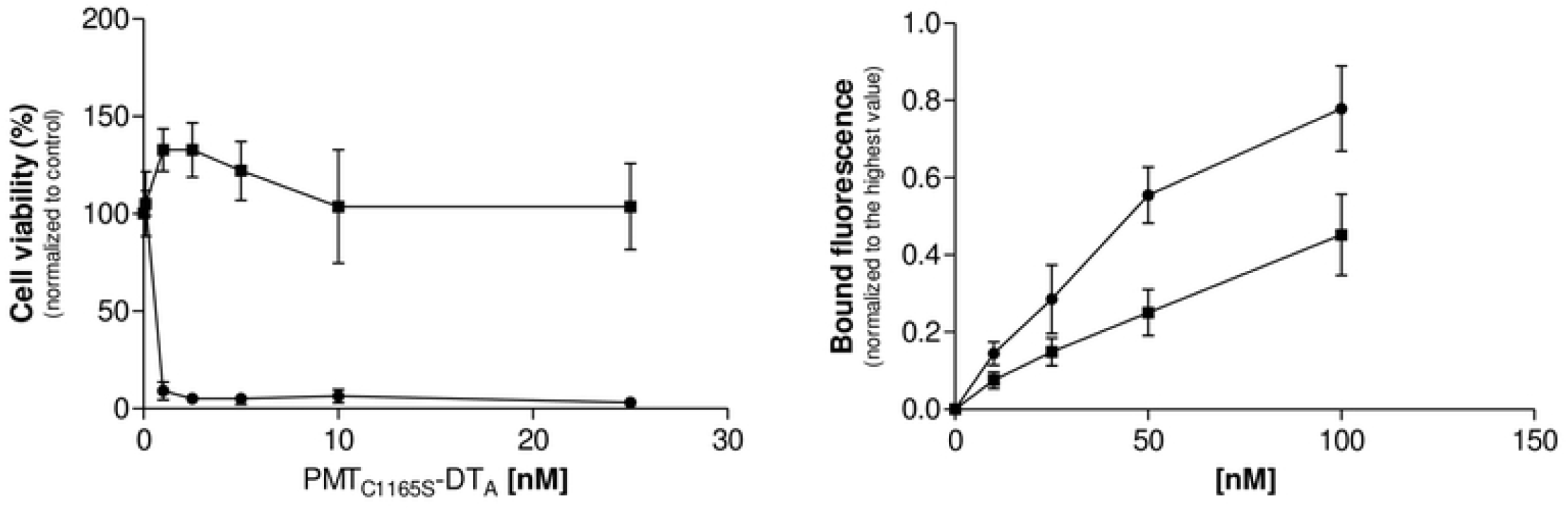
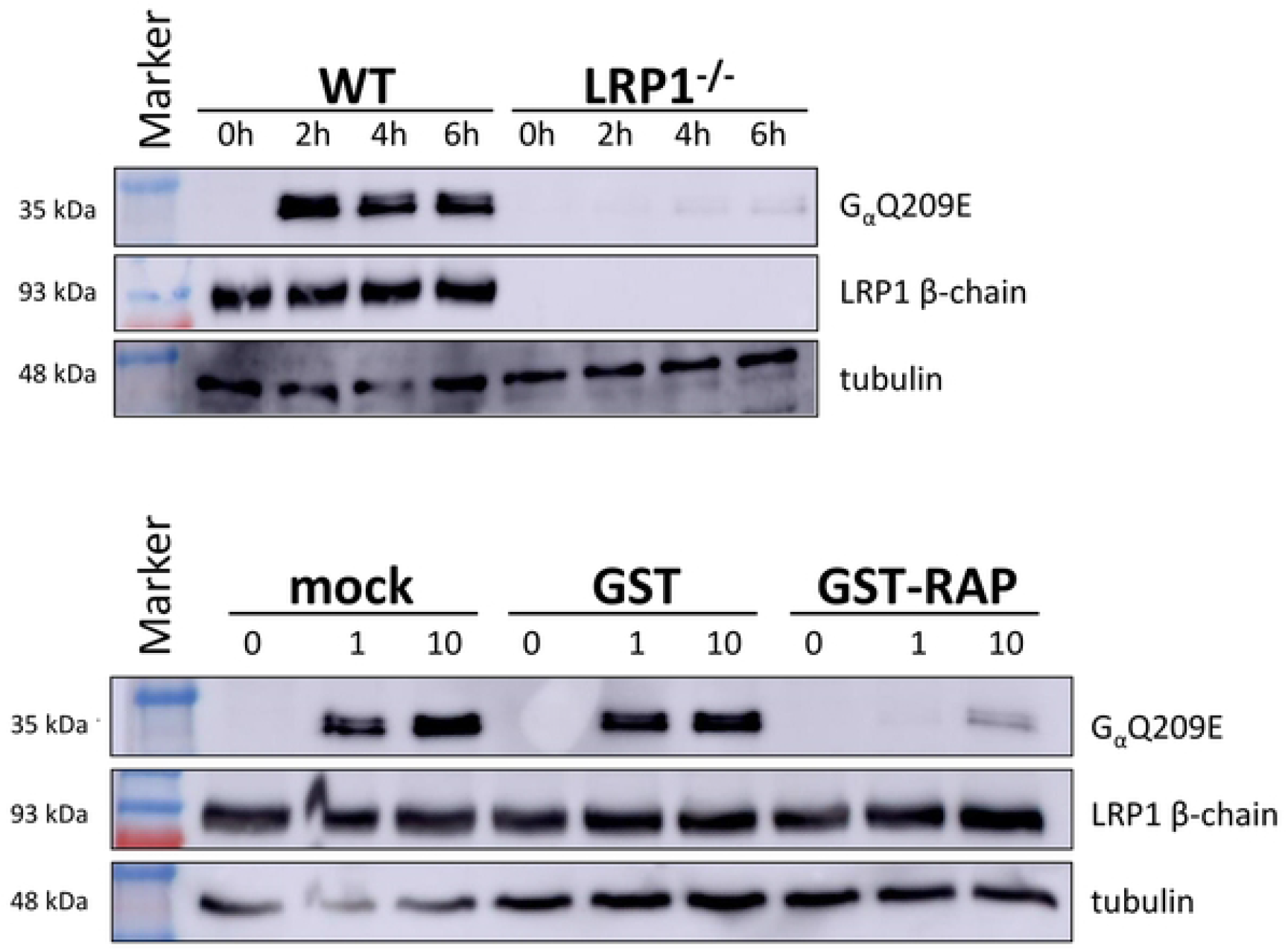
LRP1 knockout prohibits binding and uptake of PMT. MEF (circles) and MEF-LRP1^-/-^ cells (squares), respectively were incubated with increasing concentrations of PMT(C1165S)DTa for 48h and cell viability was analyzed as % of untreated controls. Shown is the mean of three independent experiments (a). Cells were incubated with increasing concentrations of Alexa488 labeled PMT (PMT_488_), washed and bound fluorescence analyzed by FACS shown as mean of three independent experiments (b). MEF and MEF-LRP1^-/-^ cells were incubated in the presence of 5nM PMT for the indicated time intervals, washed and lysed. Lysates were analyzed for toxin-induced modification of Gαq by an antibody detecting GαQ209E, for expression of LRP1 and for tubulin as loading control (c, top). MEF and MEF-LRP1^-/-^ cells were incubated in the presence of 1 or 10 nM PMT in the presence or absence of GST or GST-RAP (1 µM), respectively, washed and lysed. Lysates were analyzed for toxin-induced modification of Gαq, for expression of LRP1 and for tubulin as loading control (c, bottom).

### Direct interaction of LRP1 and PMT

To study a potential direct interaction between LRP1 and PMT, we performed a solid-phase binding assay with recombinantly produced PMT and the commercially available extracellular domains (cluster 2-4) of LRP1, respectively. These data reveal that while cluster 3 was indispensable for the PMT-LRP1 interaction PMT strongly bound to cluster 2 and 4, respectively (Fig.4). In our experiments, cluster 4 bound 18-fold stronger to PMT with a *K*_*D*_ of 2.278 nM in comparison to cluster 2 with a *K*_*D*_ of 41,65 nM.

**Fig. 4:**
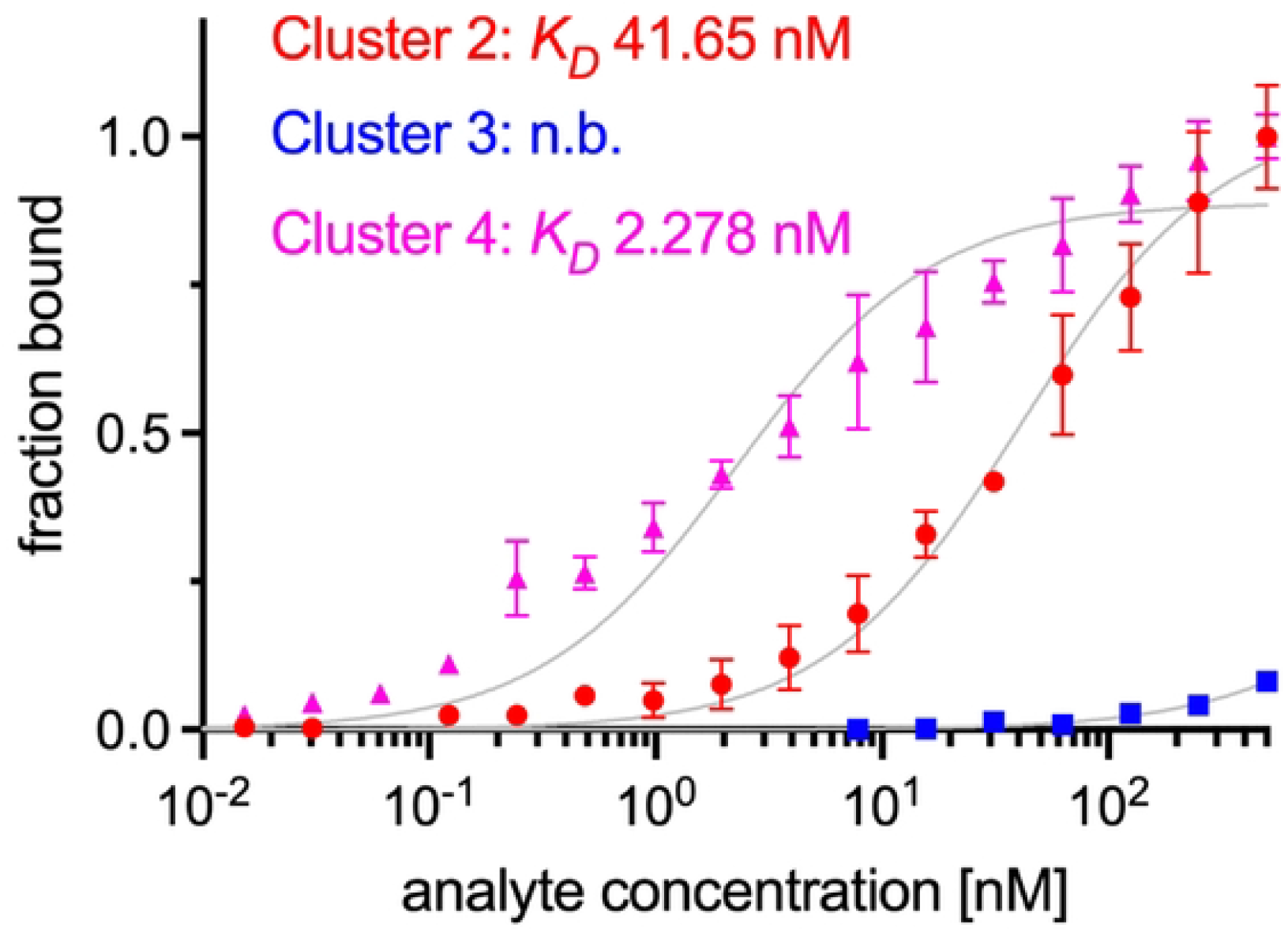
Solid-phase binding assay of PMT to clusters 2, 3, and 4 of LRP1. Graph depicts the binding curves of cluster 2 (dots), 3 (squares), and 4 (triangles) to coated His-PMT. The equilibrium-binding affinity (*K*_*D*_) is presented inside the graph. Data are shown as mean with STDEV.

### Cluster 4 of LRP1 is sufficient for PMT binding

Affinities of recombinant proteins indicate that it is mainly cluster 2 and 4 of LRP1, respectively which mediate binding of PMT with the highest affinity measured between cluster 4 and the toxin. For most viruses and bacterial toxins which interact with LRP1, the membrane closest domain (cluster 4) has been shown to be sufficient for binding and endocytosis (13). Therefore, we intended to rescue intoxication of LRP1^-/-^ cells with full-length LRP1 and with cluster 4 of LRP1, respectively. To this end, we transfected the knockout cells with plasmids encoding for the respective proteins or with the empty vector and incubated them with PMT. Whereas in cells transfected with the empty vector Gαq is not deamidated upon treatment with PMT, the protein is modified in cells re-expressing LRP1 with the maximal amount of deamidated Gαq following 6 h of treatment (Fig. 5, top). Similar results are received when we re-expressed exclusively cluster 4 of LRP1 together with the transmembrane beta chain, to ensure that the protein is localized correctly in the plasma membrane. In cells expressing cluster 4 of LRP1, Gαq is deamidated upon treatment with PMT with the maximal amount of deamidated Gα after 6 h of treatment (Fig. 5, bottom). The data show that ectopic expression of LRP1 cluster 4 restores intoxication of LRP1 knockout cells in the same manner as re-expression of LRP1 does and therefore suggest that cluster 4 sufficiently mediates binding and uptake of PMT.

**Fig.5:**
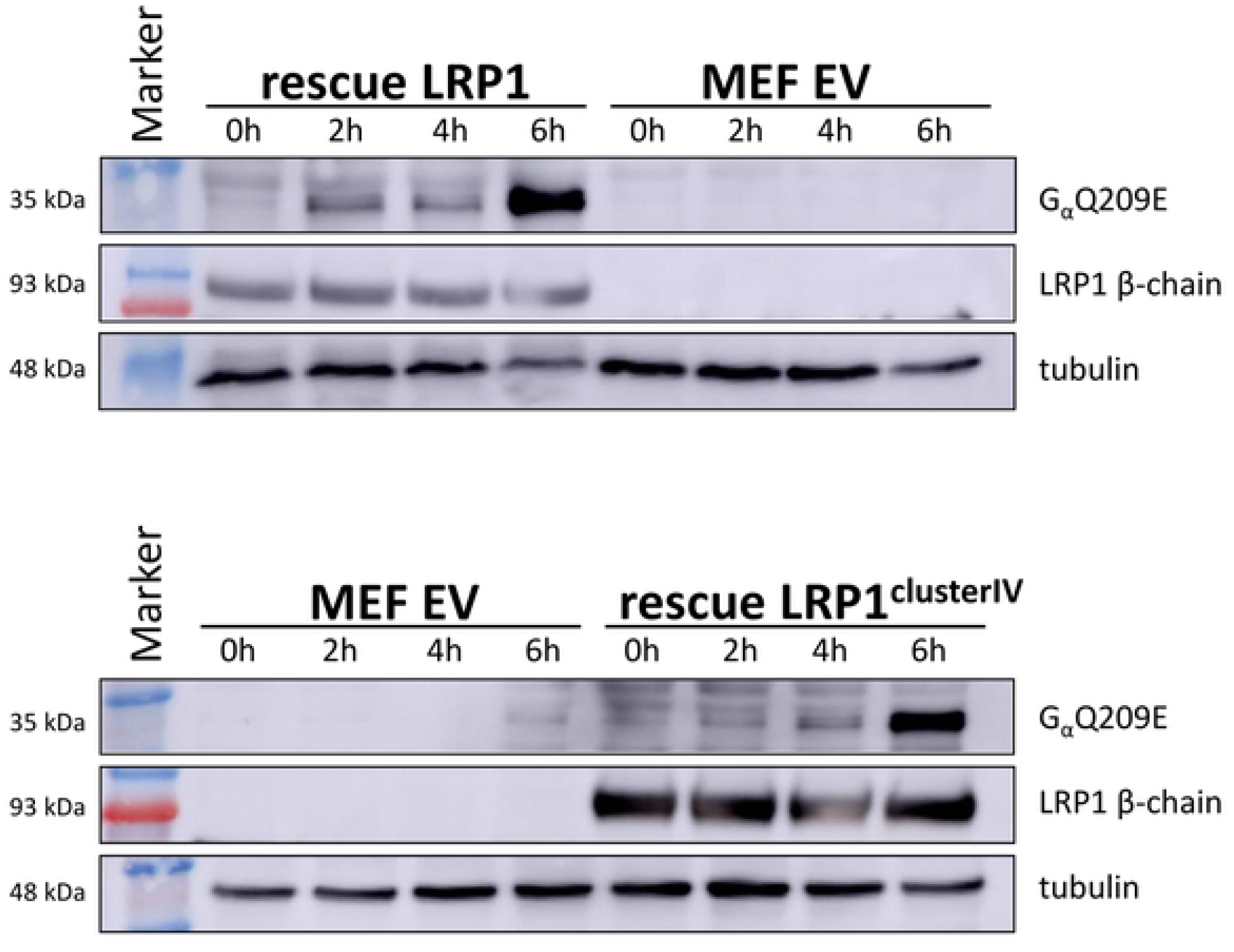
Expression of LRP1 or LRP1 cluster 4 in MEF-LRP1^-/-^ cells rescues intoxication by PMT. MEF-LRP1^-/-^ cells were transduced with retroviruses encoding LRP1 (a) or LRP1-cluster 4 (b), incubated in the presence of 5 nM PMT for the indicated time intervals, washed and lysed. Lysates were analyzed for toxin-induced modification of GαQ, for expression of LRP1 and for tubulin as loading control.

### Secretory but not epithelial cells are the main entry site of PMT into lung tissue

Following the human protein atlas (proteinatlas.org), LRP1 shows high expression in the respiratory system. The airway epithelium consists of ciliated epithelial cells, basal cells and secretory cells. We asked which of these cells preferentially take up PMT. Therefore, we made use of primary human airway tissue preparations (MucilAir™,Epithelix) which we incubated with PMT_488_ at 37°C for different time intervals, washed and fixed the tissues. Confocal optical slices were imaged to get detailed 3D information on the localization of stained PMT in the tissue. Surprisingly, the toxin appeared not equally distributed within all cells of the airway epithelium but was enriched in only a few cells (Fig. 6, z-stack shown in supplemental video (S3)). The morphology of the toxin loaded cells (flask like rounded cells without cilia) suggested that they present secretory cells. This was especially visible in orthogonal slice views of epithelial tissue preparations in which PMT localized beneath the ciliated epithelial surface of the tissue.

**Fig. 6:**
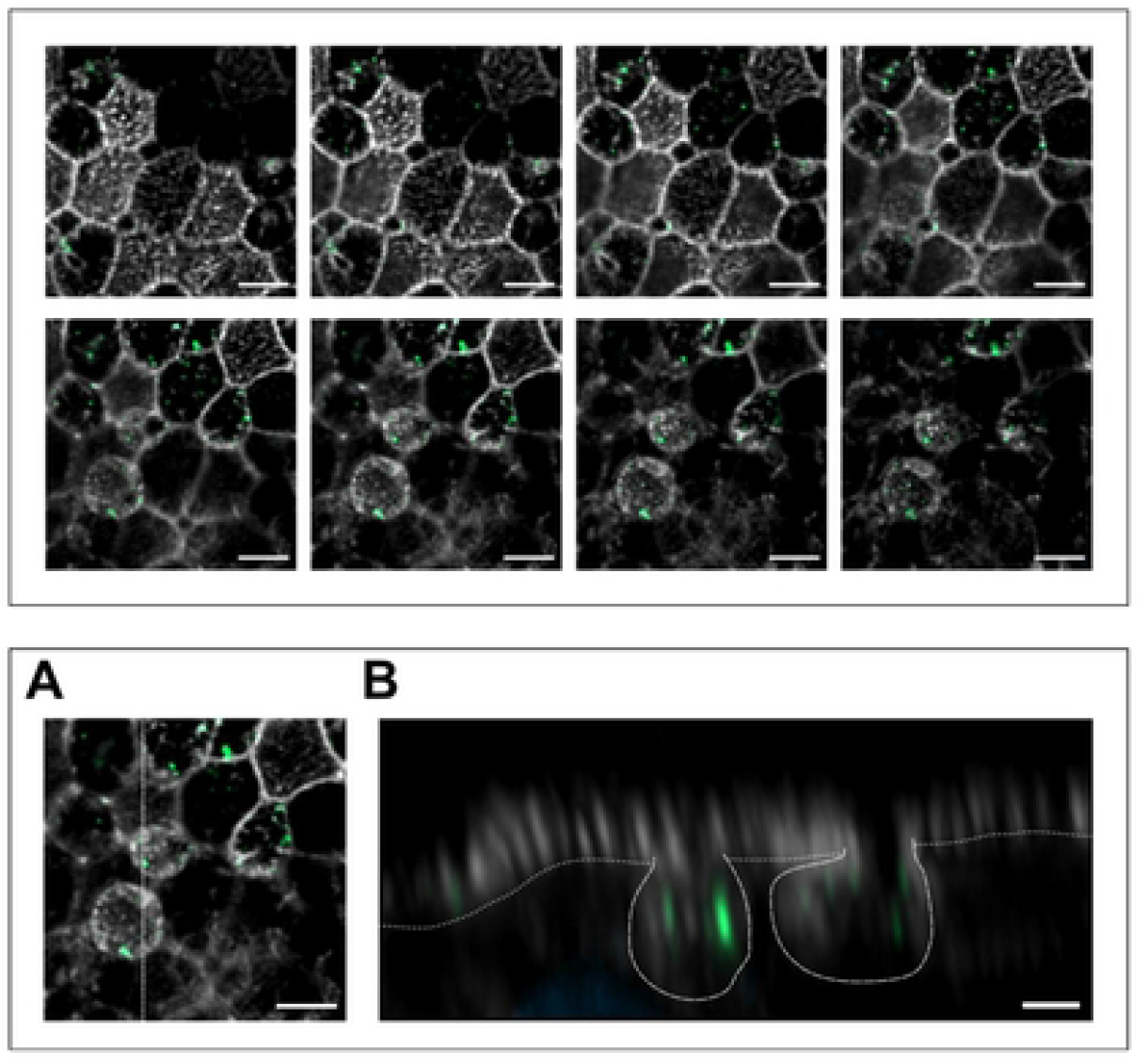
Localisation of PMT_488_ in secretory cells of human airway epithelial tissue preparations. After 4h of incubation with PMT_488_ (depicted in green), tissue preparations were fixed and stained with phalloidin-647 for actin (depicted in white). Top: Confocal optical slices were imaged for PMT_488_ in the tissue. Representative images shown are 0.52 μm apart from top to bottom, relative focus depth is shown in the top right of each image. PMT binding and uptake seem to accumulate in flask-shaped cells beneath the ciliated epithelial surface of the tissue, probably secretory cells. Dotted line in the last image marks where an orthogonal slice was reconstructed (shown in C). Bar: 5 μm. B) Scheme of orthogonal slice reconstruction in a z-stack of images. C) Orthogonal slice view of epithelial tissue preparations treated with PMT488. Confocal optical slices were imaged every 0.13 μm. Dotted lines show epithelial cell surface with cilia on top, dashed lines show the typical secretory cell shape. Bar: 2 μm. Bottom:

## Discussion

The aim of this study was to identify the cell surface receptor/s for the bacterial toxin PMT. The toxin is the main virulence factor of *Pasteurella multocida*. The non-motile, gram-negative bacterium causes diseases in animals like the atrophic rhinitis in swine. This disease is accompanied by destruction of the turbinate bone and general growth retardation (3, 29). Osteoblasts and osteoclasts seem to be main target cells of PMT. The toxin stimulates osteoclasts directly and on the other hand represses osteoblast differentiation (4). Moreover, osteocytes are activated which stimulate the osteoclastogenesis by increased secretion of the receptor activator of nuclear factor κappa B ligand (RANKL) (5)).

Using the CRISPR screen established, we identified LRP1 as top hit of the genes which when knocked out block intoxication by the PMT-DTa chimera. In line with the role of LRP1 in PMT pathophysiology of atrophic rhinitis, LRP1 signaling itself has been suggested to be involved in the differentiation of bone marrow derived macrophages into osteoclasts (30). Moreover, LRP1 on the surface of osteoblasts regulates osteoblast-osteoclast interaction and RANKL signaling (23). Both LRP1 functions may support toxin induced bone degradation. However, PMT induced LRP1 signaling has not been analyzed so far.

*Pasteurella multocida* is also found in oral secretions of dogs and cats and can cause wound infections, abscesses and osteomyelitis in man following animal bites. In immuno-compromised humans also respiratory infections with pneumonia have been described (31). In fact, LRP1 is expressed in many tissues with the highest score in the respiratory system.

A variety of ligands have been shown to interact with LRP1 including apolipoproteins, viruses and bacterial toxins with no common binding motif identified in all these ligands. Thus, LRP1 is regarded as general endocytosis receptor and moreover, LRP1 mediates intracellular signaling (32). The receptor associated protein (RAP) acts as a chaperone during synthesis of LRP1 (33) and binds to the cell surface protein when added to culture medium. Therefore, the protein acts as a competitive inhibitor for ligands binding to LRP1 (13, 34). Moreover, RAP has been used to affinity-purify LRP1 from tissues (35). In our experiments RAP competes with PMT for binding to LRP1 on cells. Since ligands for LRP1 include intracellular proteins like the nuclear protein high mobility group protein B1 (HMGB1) which is passively released upon injury, it was suggested that LRP1 has the additional function to remove such debris promoting wound healing (35). Moreover, LRP1 seems to be involved in transcytosis and may mediate uptake of bacterial toxins and viruses through the epithelium into several deeper tissues. Our results do not exclude the possibility of transcytosis since a portion of the toxin which is endocytosed for transport through the cells may reach the cytosol and therefore would be able to deamidate Gα. However, during an infection, lower toxin concentrations are expected compared to our *in vitro* conditions. LRP1 is highly expressed in the respiratory system, one of the main entry sites of *Pasteurella multocida* in animals (36). The airway epithelium consists of several cells. Interestingly, PMT was not equally distributed throughout the different cell types but highly enriched in secretory cells. The most abundant of secretory cells in the airway is the mucin producing Goblet cell. It forms Goblet cell-associated antigen passages (GAP) which mediate the transport of cargos through the epithelial gut tissue (28) for presentation of antigens to lamina propria phagocytes (37). This function seems to be independent of mucus secretion which is the established function of goblet cells in the gut and in the airway system.

However, co-staining sections from RAP_488_-treated tissues with anti-mucin antibodies negates our assumption that goblet cells express high amounts of LRP1 and therefore, preferentially may take up PMT (Fig. S4). Which of the diverse secretory and neuroendocrine cells do express LRP1 and take up PMT preferentially, remains to be determined.

Binding kinetics using a solid-phase binding assay analyzing the interaction between PMT and cluster 2, 3, and 4, respectively revealed that the toxin strongly binds to cluster 2 (*K*_*D*_ 41,65 nM) and 4 (*K*_*D*_ 2.278 nM), while cluster 3 is not required for PMT binding. The affinity of the interaction between cluster 4 of LRP1 with recombinant PMT is comparable to TpeL (KD: ∼23,2 nM) and Pseudomonas exotoxin A (KD: ∼5,2 nM) (13). The data are in line with cluster 4 presenting the site most frequently identified as interaction surface between LRP1 and diverse ligands. Moreover, cluster 4 is sufficient to bind and take up PMT into mammalian cells. Cluster 4 representing the main binding partner for cargos is a little unexpected nevertheless, because it is the one located proximate to the plasma membrane.

Besides LRP1, two genes (Dph 1 and Dph 5) encoding enzymes required for diphthamide biosynthesis were present within the first ten hits of our screen. Diphtheria toxin selectively modifies a specific diphthamide of elongation factor 2. The absence of diphthamide therefore blocks modification of EF2 and survival, proving the reasonable readout of our CRISPR screen. As control we used CAS9 expressing cells transduced with the CRISPR library but not treated with the toxin chimera. Therefore, the cells were collected earlier to use the same cell number for amplification of inserted sequences. Due to the cultivation time differences, we also expected enrichment of genes which influence proliferation. However, probably due to the strong selection force of the deadly toxin chimera, such genes were not found to be significantly enriched or diminished. In contrast, loss of tubulin binding proteins like Mical2 and Clip1 may be involved in endocytosis of PMT-DTa.

Taken together, our data show that PMT enters mammalian cells through binding and endocytosis mediated by LRP1 and that secretory cells and not epithelial cells are the main entry ports of PMT into the airway epithelium. Therefore, LRP1 may act as a common pharmacological target to interfere with uptake of disease-inducing viruses and bacterial toxins.

## Acknowledgements

We thank Silke Ludigkeit, Jürgen Dumbach and Ute Christoph for their excellent technical assistance. Moreover, we are grateful for the support by Yasuhiko Horiguchi, Osaka, who provided the plasmid encoding CAS9 and Joachim Herz, Dallas, who provided the LRP1 knockout cell line. Moreover, we thank Harald Genth, Hannover, who shared the cluster 4 of LRP1 with us. The authors acknowledge the support of the Freiburg Galaxy Team: Pavan Videm and Björn Grüning, Bioinformatics, University of Freiburg.

## Authors contribution

## Figure Legends

Fig. S1: **schematic presentation of the toxins used**

a: The Pasteurella multocida toxin (PMT) is composed of 5 domains: R: receptor binding domain, T: translocation domain, necessary for insertion into the endosomal membrane, C1-3: catalytic domain. C3 encodes for the deamidase domain. Mutation of C1165 to serine leads to a catalytically inactive toxin.

b: Diphtheria toxin (DT) is composed of three domains: DTa is the catalytic domain, T: translocation domain, R: receptor binding domain

c: The fusion protein PMT(C1165S)DT_a_ is composed of the catalytic inactive mutant of PMT and a c-terminally added catalytic domain of diphtheria toxin (DTa).

Tab. S2: table depicting the 10 mostly enriched sgRNAs and the function of the targeted genes.

Fig.S3: Non-ciliated cells take up PMT. Z-stack from Fig. 6 shown as video

Fig. S4: Localisation of RAP-GST488 in human airway epithelial tissue preparations. After 1h of incubation with 50 μM RAP-GST488 (left, green in merged image), tissue preparations were fixed and stained with phalloidin-647 for actin (depicted in white) and probed with an antibody against MUC5AC (middle, red in merged image). MUC5AC is a marker protein for goblet cells in airway tissue. RAP is a selective inhibitor and therefore binding partner of LRP1. RAP binding and uptake seem to accumulate in round, probably secretory cells, but not goblet cells. Bar: 5 μm

## Materials and Methods

### Materials

Unmodified DNA oligonucleotides were obtained from biomers.net GmbH, while HPLC grade oligonucleotides from Eurofins Genomics were used for next generation sequencing. All other reagents were of analytical grade and purchased from different commercial sources. The antibodies used in this study are designated below S1.

### Cell culture, media and growth conditions

HEK293T and Swiss3T3 were originally obtained from ATCC. MEF and MEF LRP1-/- were kindly provided by J. Herz (UT Southwestern, Dallas) and were already used in previous work of the lab. MEF EV, MEF LRP1-/-, MEF Rescue FL and MEF Rescue C4 were generated for previous work by B. Schorch (Universitätsklinikum Freiburg, Germany) and were already present at the lab. Cells were cultured in in Dulbecco’s modified Eagle’s medium (DMEM) supplemented with 10% FCS, 1% P/S and 1% NEAA. The atmosphere of 5% CO2 at 37°C was kept constantly in appropriate incubators.

### Cell lysis

For generating lysates cells were treated with RIPA buffer (1 mM EDTA, 25 mM Tris, 150 mM NaCl, 1% (v/v) Triton X-100, pH 7.4), containing “Complete” protease inhibitor cocktail (Roche) for 15 min on ice with occasional mixing. Lysates were then centrifuged at 4°C (10,000 rpm for 10 min).

Cloning, Mutagenesis and purification of recombinant PMT and PMT(C1165S)DTa PMT and PMT(C1165S)DTa were expressed as N-terminal His6-tagged proteins and purified by affinity chromatography via a Ni-NTA column as described previously (38). Cas9-expressing MEF and Swiss3T3 cells

For generation of stable cell lines expressing Cas9 the piggyBac-transposon-transposase system was used. The plasmids carrying the transposase and the transposon (pCMV-hyPBase and pPB-pgkNeo-CBh-hSpCas9n-EGFP) were gained and generated by Y. Horiguchi (Research institute of microbial diseases, Osaka). Cells were transfected by using Lipofectamine 2000 transfection reagent according to the manufacturer’s instructions with subsequent antibiotic selection with G418. Surviving cells were isolated via limiting dilution method to obtain independent clones. The clone with the highest GFP-signal was identified by using fluorescence microscopy an was further used as monoclonal origin of MEF-Cas9nEGFP and Swiss3T3-Cas9nEGFP.

### Genome-wide CRISPR/Cas9-Knock out library and screening

Generation of genome-wide Cas9-sgRNA library and subsequent screening were performed as described by Horiguchi et al., 2020 and adapted. We utilized a well-functioning and validate genome-wide mouse lentiviral CRISPR guide RNA library v2 (Addgene). It consists of 90230 gRNA sequences directed against 18424 different genes across the mouse genome. HEK293T cells were co-transfected with library plasmids and suitable packaging- and envelope plasmids (CRISPR & MISSION® Lentiviral packaging mix, Sigma-Aldrich) by using Lipofectamine 2000 Transfection Reagent according to the manufacturer’s instructions. Lentiviral particles were obtained after 48 h via centrifugation of the culture supernatant at 1500 × g for 15 min at 4°C and subsequent sterile filtration trough 0,45µm pore filters. The virus titer was quantified by an ELISA assay against the lentiviral envelope protein p24 (HIV1 p24 ELISA Kit, Abcam). The Cas9 expressing MEF cells were infected in presence of 8 µg/ml Polybrene with the lentiviral particles at a multiplicity of infection of 0,3 to avoid multiple gene knock outs in one cell. Following the infected cells were identified by antibiotic selection procedure by using 8 µg/ml Puromycin after 48 h.

For screening the library cells were incubated 3 times with 2µg/ml PMT-DTa for 36 h with an intermediate interval of 24 h in toxin-free full medium. The surviving cells were cultivated to confluence of a 6 well and the genomic DNA was isolated by incubation of the harvested cells with proteinase K at 37°C in accordance to instructions. The sgRNA containing sequences of the genomic template DNA were amplified by PCR using the indicated Primers of HPLC grade (S2) and Q5 Hot Start high-fidelity DNA polymerase (New England BioLabs). The PCR products of the toxin resistant cells and untreated library cells were analyzed by deep sequencing using an Illumina-based procedure via a commercial supplier (Eurofins Genomics).

### Cell viability assay

The metabolic activity of MEF and MEF LRP1-/- cells (cell viability) was determined by using the CellTiter-Blue® cell viability assay (Promega) following the manufacturer’s protocol. The fluorescent product resofurin was measured on a multimode microplate reader (Infinite M200; Tecan).

### Immunoblotting

For immunoblotting cell lysates were subjected to SDS-PAGE and electro transferred onto a PVDF membrane. After subsequent blocking and washing, PMT action was detected by a deamidation specific antibody anti-Gαq Q209E as described before (39). LRP1 and tubulin were visualized by suitable antibodies given in S1. By utilizing SignalFire enhanced chemiluminescent detection reagent (Cell Signaling Technology) and the imaging system LAS-3000 (Fujifilm), binding of the compatible horseradish peroxidase-coupled secondary antibody was detected.

### Solid-phase binding assay

N-terminally His-tagged Pasteurella multocida toxin (PMT) was coated at 10 µg/mL (500 ng/well) onto 96-well plates (NuncMaxisorb) at 4°C overnight. After washing three times with TBS supplemented 2 mM CaCl_2_, plates were blocked with Pierce™ protein-free TBS blocking solution (pH 7.4; Thermo Fisher Scientific) at room temperature (RT) for 2 h. Ligands (LRP1 cluster 2, 3, and 4) were diluted to concentrations from 0.02 nM to 500 nM and incubated at RT for 2 h. After extensive washing with TBS supplemented 2 mM CaCl_2_, bound ligands were detected using a goat anti-human IgG-HRP-conjugated antibody (Santa Cruz, sc-2453; 1:4000). HRP was detected by Pierce TMB ELISA Substrate (Thermo Fisher Scientific). Absorption was measured at 450 nm after stopping the reaction with 2 M sulfuric acid. A blank value corresponding to BSA blocked wells of the respective analyte concentration was automatically subtracted.

### FACS analysis

For binding studies PMT was labelled covalently with DyLight488 maleimide (Thermo Fisher Scientific) according to the manufacturer‘s instructions. Pre-cultured MEF cells and derivates were detached from culture plates using 10 mM EDTA in PBS. For each condition 2 × 105-2.5 × 105 cells were used and incubated for 20 min with the indicated PMT-488 concentration at 4°C. Following several washing steps with ice-cold PBS, cell bound fluorescence was measured using a FACS Melody (BD Bioscience). Via the appropriate Software FACSChorus™ (BD Bioscience) 10.000 cells were analyzed. After gating and the average of surface bound fluorescence of all cells was given.

### Immunofluorescence

Treated airway epithelial tissue preparations (MucilAir) were fixed using 4% paraformaldehyde in PBS (pH 7.4) for 15-20 min, washed three times with PBS and permeabilized with 0.25% Triton X100 for 10 min. Primary antibodies against Muc5ac were incubated overnight at 4° C. Secondary antibodies anti-rabbit AlexaFluor-568 (Invitrogen #A11011; 1:500) or anti-mouse AlexaFluor-568 (Invitrogen #A10037; 1:500) were incubated together with ATTO-647 coupled phalloidin (Hypermol #8817-01; 1:400) for 1h at room temperature in the dark. The airway epithelial tissues were mounted using ProLong Glass with DAPI (Invitrogen) for staining of nuclei and studied on a Zeiss LSM800 with Airyscan acquisition using a 63x/1.4 NA oil objective. Images were analysed and 3D rendered using ZEN blue software (version 3.2, Zeiss) and ImageJ.

Immunofluorescence of cryosections: after the fixation step the airway epithelial tissues were cryoprotected in 30% sucrose/PBS at 4° C overnight. Airway tissues were transferred to plastic cryomolds and embedded in OCT compound (Tissue Tek, Sakura) and frozen on an aluminium block sitting in liquid nitrogen. 9μm cryosections were obtained on a Leica cryostat (CM 1950) and melted onto plus charged microscope slides, air dried and stored at −20°C. After rehydration (10-20 min in PBS at RT), cryosections were stained and imaged as epithelial tissue samples (see above).

### Statistics

Statistical analysis and presentation were performed using the software GraphPadPrism 5 (GraphPad, San Diego, USA). Unless otherwise indicated, the mean value and the standard deviation the mean value and the calculated standard deviation (SD) are shown in all diagrams, unless otherwise stated, SD), are shown. P-values smaller than 0.5 were defined as statistically significant and marked with an asterisk (n.s. = not significant; *, p < 0.05; **, p < 0.01; ***, p < 0.001; ****, p < 0.0001).

**S1.** Antibodies used in this study

S1 Antibodies used in this study
| Antibody | Source |
| --- | --- |
| GαQ209E | self-produced in hybridoma cells (39) |
| LRP1 | Abcam |
| Tubulin | Santa Cruz |
| Anti-rat IgG HRP | Cell Signaling |
| Anti-goat IgG HRP | Cell Signaling |
| Anti-mouse IgG HRP | Rockland Inc. |
| Anti Muc5ac | Thermo Fisher Scientific |

**S2.**
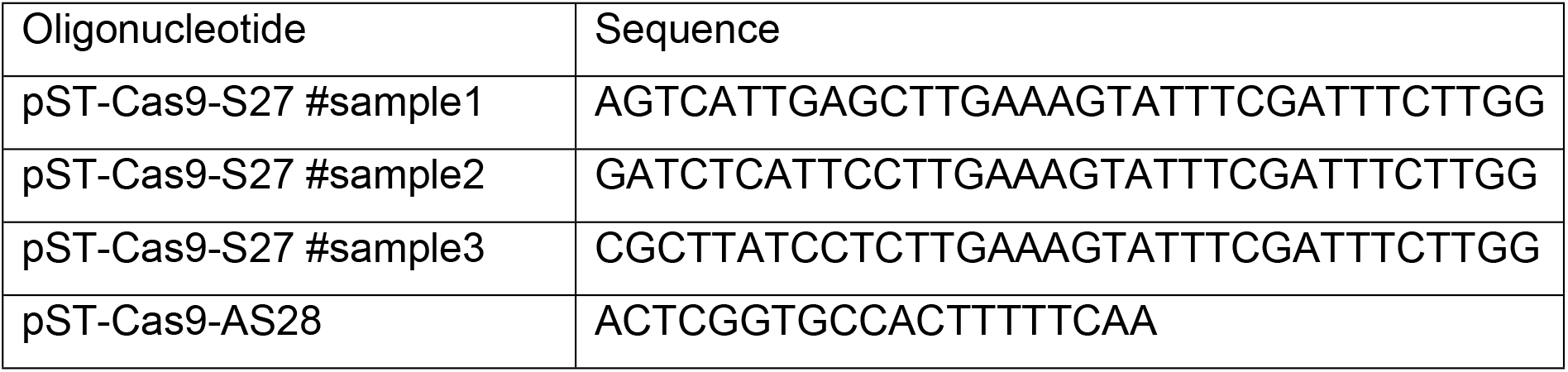
Oligonucelotides used in this study

### CRISPR data analysis

The gene expression changes analysis was performed using the three treated and two control single-end sequencing libraries. The sequencing quality was assessed using FastQC and MuliQC (40, 41). All read libraries were mapped against an sgRNA FASTA file, containing the full sgRNA library sequences, using BWA-MEM with an adjusted minimum score output of 18 and enabling penalties for 5’-end and 3’-end clipping (42, 43). Mapping rates were analyses using Samtools stats (44), resulting in an average rate 65%. The quantification was done using featureCounts (45). The expression changes between the two conditions were calculated with DESeq2 (46). The gene expression changes were summarized in a volcano plot using p-value 0.05 and LogFC 0.584 thresholds (47). The full data analysis was performed in Galaxy, see: https://usegalaxy.eu/u/teresa-m/h/crisprsgrnadgeanalysis.

